# Lead Exposure During Differentiation Perturbs Electrophysiological Function in iPSC-Derived Cardiomyocytes in a Cell Line-Dependent Manner

**DOI:** 10.64898/2026.10.01.755665

**Authors:** Kimberley E. Sala-Hamrick, Esha K. Shah, Andre Monteiro Da Rocha, Laurie. K. Svoboda

## Abstract

Lead (Pb) exposure is strongly associated with cardiovascular diseases, but the effects of developmental Pb exposure on the electrophysiological properties of human cardiomyocytes have not yet been investigated. Using human induced pluripotent stem cell (iPSC)-derived cardiomyocytes, we investigated the effects of human-relevant Pb exposure during ventricular cardiomyocyte differentiation on action potential properties and calcium dynamics in differentiated myocytes. We exposed 2 iPSC lines, from 1 male and 1 female donor, to Pb (0.5 µM or 5 µM) or control during only the first 8 days of differentiation, prior to functional maturation. Differentiation was carried out using an established protocol involving temporal modulation of Wnt signaling, followed by purification and maturation of contractile myocytes. We measured action potential properties and calcium dynamics using fluorescent dyes and optical mapping. We hypothesized that Pb exposure would adversely affect cardiomyocyte function, with effects persisting after cessation of exposure. In keeping with this hypothesis, we found significant Pb-induced changes in calcium (Ca) transients and action potential dynamics that were concentration and cell line specific and persisted well after exposure had ended. In the female cell line, beat frequency significantly increased with Pb exposure for both action potentials and Ca transients, while in the male cell line, beat frequency significantly decreased with Pb exposure for action potentials and increased for Ca transients. For voltage measurements, we observed significantly increased arrythmias with Pb exposure in both cell lines. Significant changes in repolarization and recovery time (for voltage and Ca, respectively) were present with Pb exposure in both cell lines. Gene expression of Ca transporters/channels and K channels were not affected, suggesting that persistent effects of Pb exposure on cardiomyocyte electrophysiology may occur through post-translational mechanisms. Overall, this study paves the way for future work to examine how exposure to Pb impacts long-term cardiac function through developmental programming.

**Statements and Declarations:** All authors declare that they have no conflicts of interest.

## Introduction

Pb, a naturally occurring element, is very toxic to humans, yet exposure to this pollutant continues to be a worldwide problem.(1) Epidemiological studies have strongly and consistently shown a relationship between Pb exposure and the risk of various cardiovascular diseases, including arrhythmias and other conduction defects.(2,3) However, how early life Pb exposure may alter cardiac electrophysiological function through developmental programming is largely unexplored. In the current study, we exposed iPSCs to Pb during the early stages of ventricular cardiomyocyte-directed differentiation and examined action potential and calcium (Ca) transient function in the resulting purified myocytes using optical mapping.

The Developmental Origins of Health and Disease (DOHaD) hypothesis posits that environmental exposures during key periods of development can lead to dysfunction or disease later in life.(4) Human and animal studies demonstrate that developmental Pb exposure has deleterious effects on heart development and cardiovascular health. Prenatal Pb exposure has been associated both with an increased risk of congenital heart defects as well as increases in blood pressure in children.(5–10) Work in rodent models has also found that prenatal Pb exposure leads to later cardiac dysfunction.(11,12)

The arrhythmogenic effects of Pb have been demonstrated in human and animal studies, and have been largely attributed to autonomic nervous system toxicity.(13) Less is known, however, about how Pb exposure during development may directly alter cardiac electrophysiological properties, independent of autonomic dysfunction.

Briefly, cardiac electrophysiology is characterized by the coupling of electrical stimulus (action potential) to an intracellular rise in Ca in cardiomyocytes that leads to the production of a rhythmic heartbeat through a process called excitation contraction coupling (ECC).(14,15) Importantly, Pb is known to mimic calcium (Ca), which is pivotal for the normal coupling between cardiac action potentials and contractility. Pb and Ca are both divalent cations of similar ionic radius, and disruption of Ca signaling is one mechanism by which Pb exposure may lead to cardiac arrhythmias through direct effects on cardiomyocytes.(16–18)

In this work, we exposed 2 human iPSC lines, from 1 male and 1 female donor, to a human exposure relevant concentration of Pb (0.5 µM) and a concentration representing acute Pb poisoning (5 µM) during the first 8 days of their differentiation to ventricular cardiomyocytes, after which cardiomyocytes were functionally matured and purified. We chose this exposure window, prior to the onset of spontaneous contraction and functional maturation of cardiomyocytes (days 8-30), to capture the developmental period where pluripotent cells first differentiate into mesodermal and then progenitor lineages.(19) To assess changes in cardiomyocyte function, we conducted optical mapping of action potentials and Ca transients in the two differentiated cardiomyocyte (iPSC-CM) lines. To determine whether the observed effects of Pb were due to altered expression of genes that underlie cardiac excitability and calcium handling (Ca handling genes *CACNA1C, RYR2, ATP2A2*, *SLC8A1,* and potassium channel genes *HERG1A/B*), we examined their expression using qRT-PCR.

We found that Pb exposure during differentiation had lasting, cell line-specific effects on cardiomyocyte electrophysiology, suggesting altered ion channel function and calcium handling. Importantly, these effects persisted well beyond cessation of exposure, suggesting potential developmental programming. Overall, our findings suggest that developmental Pb exposure has durable and lasting effects on cardiomyocyte electrophysiology, revealing an important mechanism of Pb cardiotoxicity in humans.

## Methods

### Cell culture: iPSC maintenance

The iPSC lines used in this study were purchased from WiCell (Madison, WI, USA). The iPSC lines were reprogrammed from peripheral blood mononuclear cells that were isolated from a 24-year-old Caucasian female (PENN002i-442-1) and a 21-year-old Caucasian male (PENN042i-258-12). 24-well and 6-well Corning plates were coated with Corning Matrigel Basement Membrane Matrix at a 1:40 dilution in DMEM:F12 HEPES (#07-200-80, #09-761-146, #356234, and #11330032; Thermo Fisher Scientific, Waltham, MA USA). iPSCs were cultured in feeder free conditions with StemMACS iPS-Brew XF Media or StemMACS PSC-Brew XF (#130-104-368 and #130-127-865; Miltenyi Biotec, Bergisch Gladbach, Germany) and maintained in an incubator at 37°C and 5% CO2. Media changes were performed daily for cells cultured in iPS-Brew and every other day for PSC-brew with media pre-warmed in a bead bath to 37°C. iPSCs were viewed under a microscope daily, and colonies showing signs of spontaneous differentiation were marked and aspirated. When cells reached 80% confluency, they were dissociated non-enzymatically using Versene (#15040066, Thermo Fisher Scientific). After passaging, iPSCs were re-plated at a density of 1:24 for maintenance plates, and the remaining cell stock was used to plate a monolayer (80-100% of cells-see below).

### Cell culture: iPSC differentiation

The differentiation of iPSCs into cardiomyocytes was initiated in 1-3 days post-plating, when the monolayers reached 80-95% confluence. Differentiation was carried out using a modified GiWi protocol as previously described and outlined in **Figure 1**.(20,21) Cells were maintained in basal media (RPMI with HEPES CAT# 22400-89 Gibco, Grand Island, New York, USA) with supplements ascorbic acid #A8960 and BSA #A9418 (Sigma Aldrich, St. Louis, MO, USA) for the first 8 days of the differentiation process. iPSCs were directed into a mesoderm state with activation of Wnt signaling induced by the addition of the GSK-3 inhibitor CHIR99021 (#4423) into the basal media on day 0.(19,22) After 2 days of incubation, Wnt signaling was inhibited with the addition of IWP4 (#5214, Tocris Bioscience, Avonmouth, Bristol, United Kingdom) to drive cells towards a cardiac lineage.(19,22) On days 4 and 6, media changes were performed using basal media with no additional small molecules. From day 8 until purification, B27 + insulin media was used to maintain the cells with media changes occurring every other day (RPMI # 11875-093 and B27 supplement #17504-044; Gibco).

**Figure 1:**
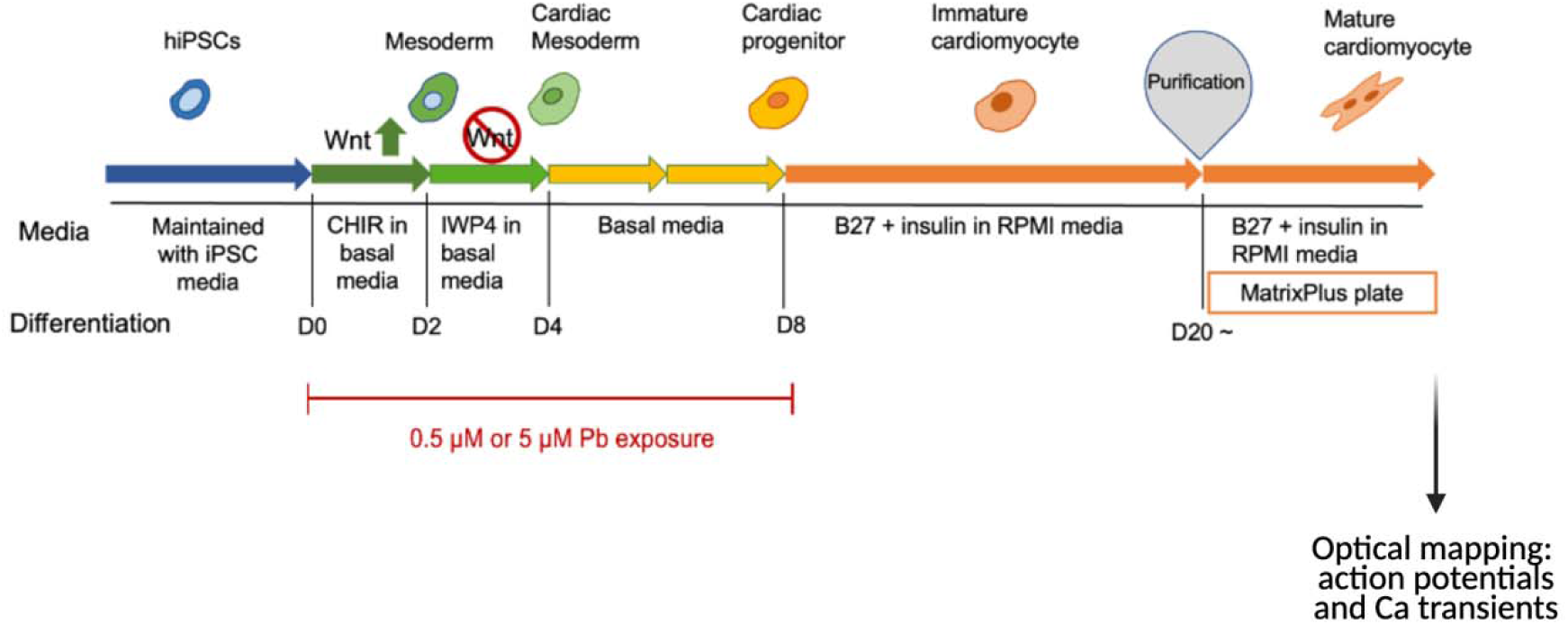
Ventricular differentiation protocol of iPSCs into cardiomyocytes and experimental design. Differentiation protocol and subsequent purification, with Pb exposure on the first 8 days of differentiation, purification between days 10-20, and optical mapping performed for purified cardiomyocytes 7-10 days after purification.

### Cell culture: Pb exposure paradigm

Pb (II) Acetate Trihydrate (CAS: 6080-56-4; #215902, Sigma-Aldrich) was diluted in tissue culture grade water. As shown in **Figure 1**, iPSCs undergoing differentiation were exposed for the first 8 days (days 0-8) at an environmentally and occupationally relevant concentration of 0.5 µM and a higher toxic concentration of 5 µM. These experiments were performed in 6 well plates with at least 2 technical replicates per condition.

### Cell culture: cardiomyocyte purification

Cardiomyocytes derived above (both control and Pb exposed) were purified once beating was visualized throughout each well (days 10-20), using the PSC-Derived Cardiomyocyte Isolation Kit (#130-110-188, Miltenyi Biotec, Bergisch Gladbach, Germany) as described.(23) This kit employs a selective depletion strategy to enhance the cardiomyocyte purity to 98%. First, cells were enzymatically dissociated with trypsin (#25200056, Thermo Fisher Scientific). After trypsin inactivation with EB20 media (reagents in **Table 1** below), cells were passed through 30 um pre-separation filters (#130-041-407 Miltenyi Biotec), centrifuged, washed with magnetic-activated cell sorting (MACS) buffer (#130-091-221, Miltenyi Biotec) centrifuged again, and then incubated in cardiomyocyte depletion primary antibody from the Miltenyi isolation kit for 10 minutes. Primary antibody was washed off with MACs before incubation with secondary antibody from the kit which attaches the non-cardiomyocyte cells to magnetic microbeads. After this, cells in MACs suspension were passed through magnetic separation columns attached to a magnet, where cardiomyocytes pass through and noncardiomyocytes remain attached to the column (LS columns #130-042-401 and magnetic separator #130-042-303, Miltenyi Biotec). Cells were then spun down and redissolved in EB20 media. Finally, the purified cardiomyocytes were plated at 75,000 cells per well on a 96 well CELLvo™ Matrix Plus extracellular matrix coated plate (#30001.57, StemBioSys, San Antonio, Texas, USA). After 48 hours, cells were then maintained in EB20.

**Table 1:** Reagents in EB20 Media.

| Reagent to make EB20 media | Product Number, Company, Location |
| --- | --- |
| 80% DMEM:F12 | #11-220-033, Gibco, Grand Island, New York, USA |
| 20% FBS | #f2-442-500ml, Sigma-Aldrich, St. Louis, MO, USA |
| 1 mmol/L L-glutamine | #25030-081, Gibco, Grand Island, New York, USA |
| 0.1 mmol/L Nonessential amino acids | #11140-050, Gibco, Grand Island, New York, USA |
| 0.1 mmol/L beta-mercaptoethanol | #21985-023, Invitrogen, Waltham, Massachusetts, USA |
| 0.001% Blebbistatin | #13186, Cayman chemical, Ann Arbor, Michigan, USA |

### Optical mapping of action potentials and Ca transients

After cells recovered from purification (beating again throughout each well and matured after 7-10 days), optical mapping of action potentials and Ca transients was performed to measure electrophysiological changes in cells exposed to Pb during differentiation (**Figure 1**). Two different fluorescent dyes were used: for action potentials, FluoVolt™ Membrane Potential Kit (#F10488 Thermo Fisher Scientific) and for Ca transients, Calbryte 520 AM (#50-224-9887 Thermo Fisher Scientific). Cardiomyocytes in a 96 well plate (purified as described above) were exposed in parallel plates to each reagent for 30 minutes and washed with HBSS++. After this, optical mapping was performed on a Nautilai plate reader within 5 minutes (Curi Bio, Seattle, WA, USA). Briefly, cells were placed in a motorized, temperature-controlled stage (37°C). Fluorescence signals were acquired using preconfigured optical settings of the Nautilai for both FluoVolt (voltage) and Calbryte (Ca) imaging, recorded for 20-30 seconds at a frame rate of 250 frames per second and LED intensity of 10% for action potentials and 5% for Ca transients.

Nautilai software purpose-built for the plate reader processed the data, and a custom developed parallel StemBioSys software was used to assess the resulting action potentials and Ca transients in each well (StemBioSys Optical Electrophysiology Analysis Tool (.oeat), StemBioSys, Inc./Southwest Research Institute).(21) Together, this allows for comprehensive analysis of a range of parameters for both action potentials and Ca transients. Both action potential and Ca transient duration data were corrected for beat frequency using Fridericia’s equation: Corrected Duration = Duration / (Cycle Length) ^ (1/3) where Cycle Length = 1 / Beat Frequency in Hz. Each optical mapping parameter was graphed in prism and significance between treatments was tested using one-way ANOVAs. To determine percent of arrhythmia occurrence, each region of interest was analyzed for non-consistent beat patterns (beyond slowing down or speeding up), and the percent of arrhythmia regions per condition per replicate was computed, graphed, and analyzed for significance using a mixed-effects binomial generalized linear mixed model to account for replicate-to-replicate variation.

### RNA extractions

Cells that had been frozen at −80 after optical mapping were scraped off plates for RNA extraction, which was performed using the *mir*Vana™ miRNA Isolation Kit (large RNA fraction; #AM1560 ThermoFisher) according to the manufacturer’s instructions with an additional DNAse incubation step (#79254, Qiagen, Aarhus, Denmark). Sample concentration and quality were assessed using a nanodrop microvolume spectrophotometer (ThermoFisher). Three replicates of purified cardiomyocytes from unique differentiations used in optical mapping were extracted and used for qRT-PCR.

### qRT-PCR

500 ng of total RNA was reverse transcribed into cDNA using the iScript cDNA Synthesis Kit (#1708890 BioRad, Hercules, California). mRNA expression of genes whose primers are listed in **Table 2** below were quantified using the CFX Real-Time PCR using iTaq Universal SYBR Green Supermix according to manufacturer’s protocols with all samples run in triplicate. mRNA expression was compared using the same method on the same plate to amplification of two reference genes (*ACTB and GAPDH*; primer information in **Table 2** below). All qRT-PCR reactions were carried out for 40 cycles under standard PCR conditions. ΔCq values were calculated by subtracting the Cq triplicate average of a target sample from that of the geometric mean of the reference genes. The relative difference in gene expression was then calculated using 2^−ΔCq^ and graphed using GraphPad Prism, where one way ANOVA and unpaired t-tests were used to determine statistical differences between control and treatment groups.

**Table 2:**
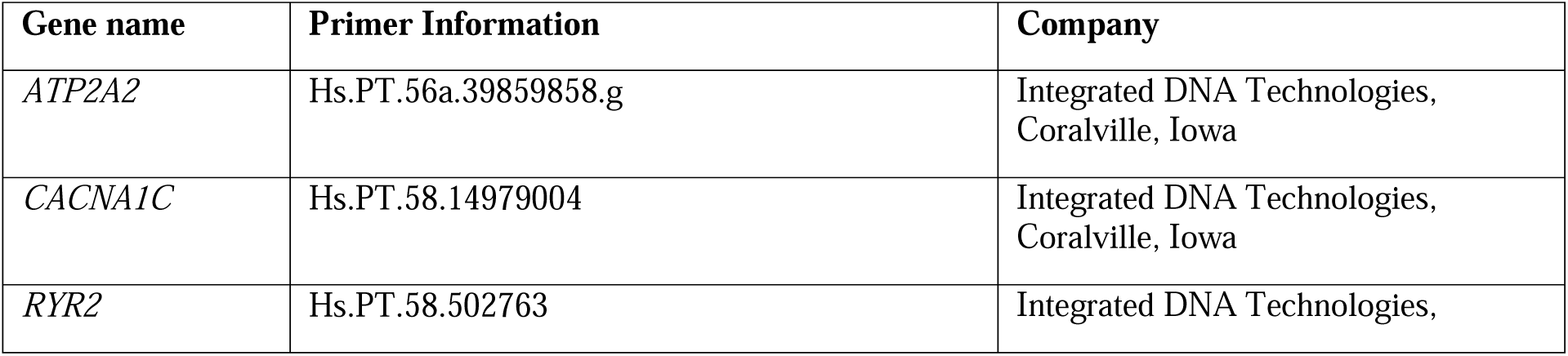
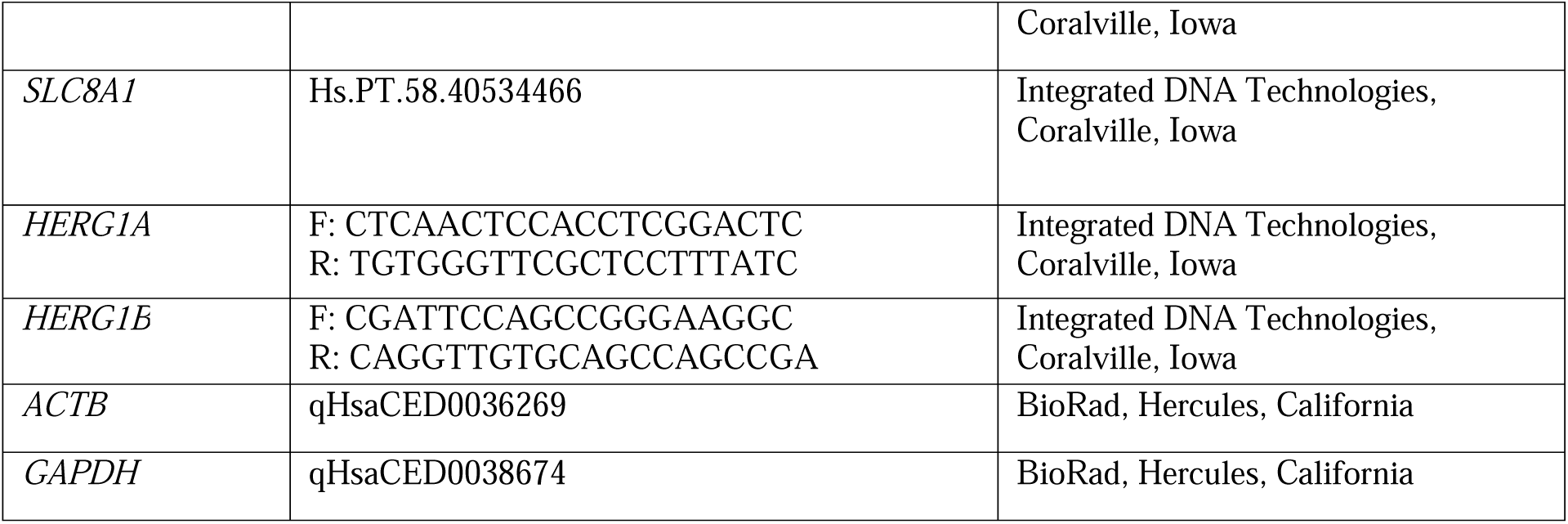
Primer Information.

## Results

To assess functional consequences of Pb exposure during differentiation on cardiomyocyte electrophysiology, we performed optical mapping of action potentials and Ca transients. Two cell lines were exposed to 0.5 or 5 µM Pb or vehicle control during the first 8 days of differentiation. The remaining protocol (maturation and purification) was then carried out in the absence of Pb exposure (**Figure 1**). Therefore, upon optical mapping, cells had not been exposed to Pb for 9-17 days.

### Exposure to Pb during differentiation affects cardiomyocyte action potentials in a cell line-specific manner

Beat frequency, or the number of action potentials per second (Hz) was significantly increased with 5 µM Pb exposure during differentiation in the female line but decreased with 0.5 µM Pb exposure in the male line (**Figure 2b**). Action potential duration at 80% repolarization (APD80), or the repolarization time, was increased in the female line at both concentrations and decreased in the male line at 0.5 µM Pb only (**Figure 2c**). Very modest but significant changes in baseline fluorescence were seen for both cell lines, with a significant increase at 5 µM in the female cell line and decrease at 0.5 µM in the male cell line (**Figure 2d**). There was also an increase in the action potential upstroke slope at 0.5 µM in the female cell line only (**Figure 2e**). Sample traces for action potentials are shown in **Figure 2a**.

**Figure 2:**
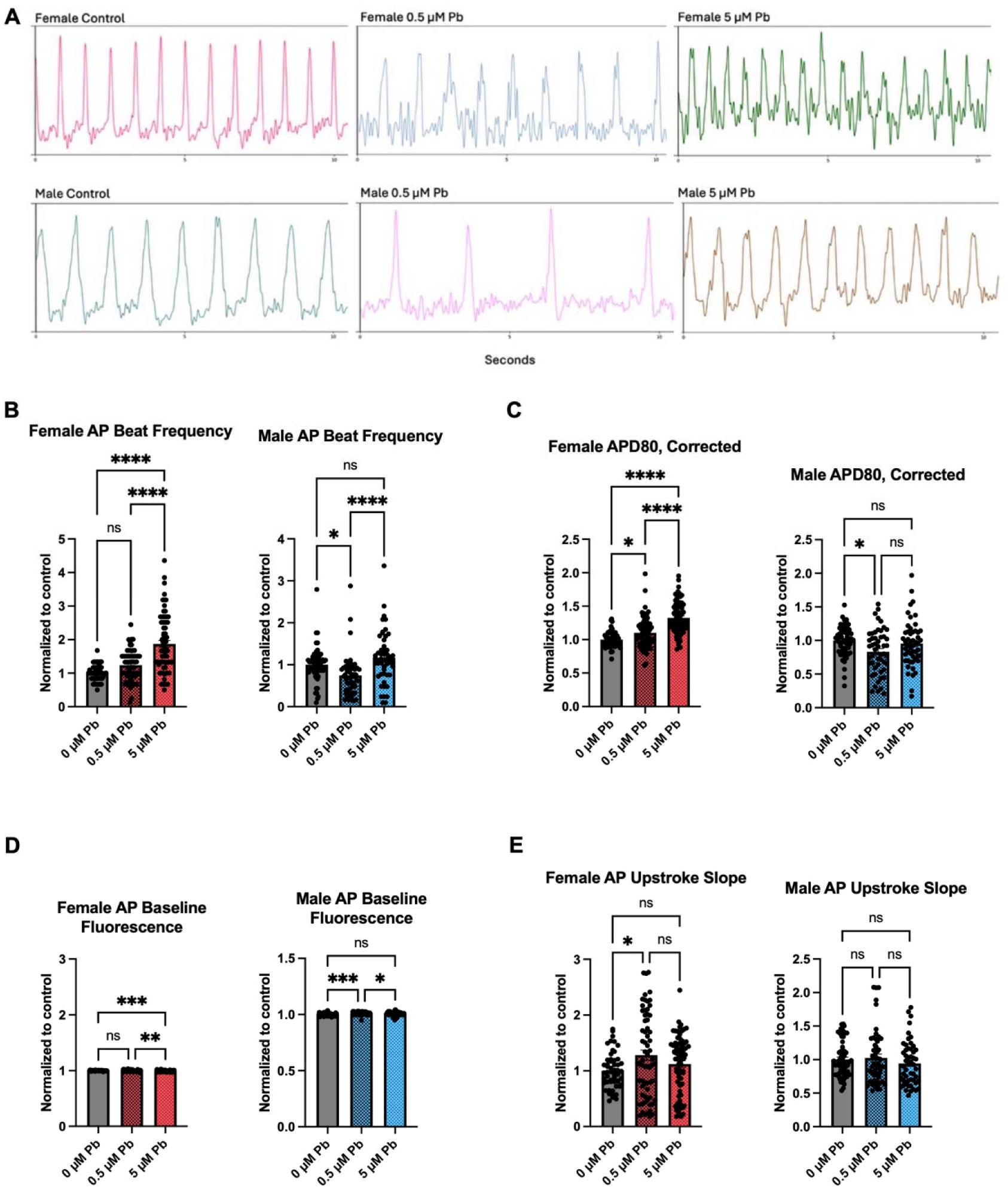
Action potential changes in female and male iPSC-derived ventricular cardiomyocytes exposed to Pb during differentiation. Sample action potential beat traces (A), significant changes in beat frequency (B), APD80 (C), and Baseline Fluorescence (D) were observed for both cell lines, and a change in upstroke slope was observed for the female cell line only (E). (n= 3 female and n=4 male replicates; one-way ANOVA; * p-value < 0.05, ** p-value < 0.01, *** p-value < 0.001, **** p-value < 0.0001).

Occurrence of arrhythmias were also assessed in the same samples that underwent optical mapping (example arrhythmic beat traces shown in **Figure 3a**), and a significant increase in percent arrhythmias as assessed by action potentials was observed for both concentrations of Pb for both the female derived and male derived iPSC-CMs (**Figure 3b**).

**Figure 3:**
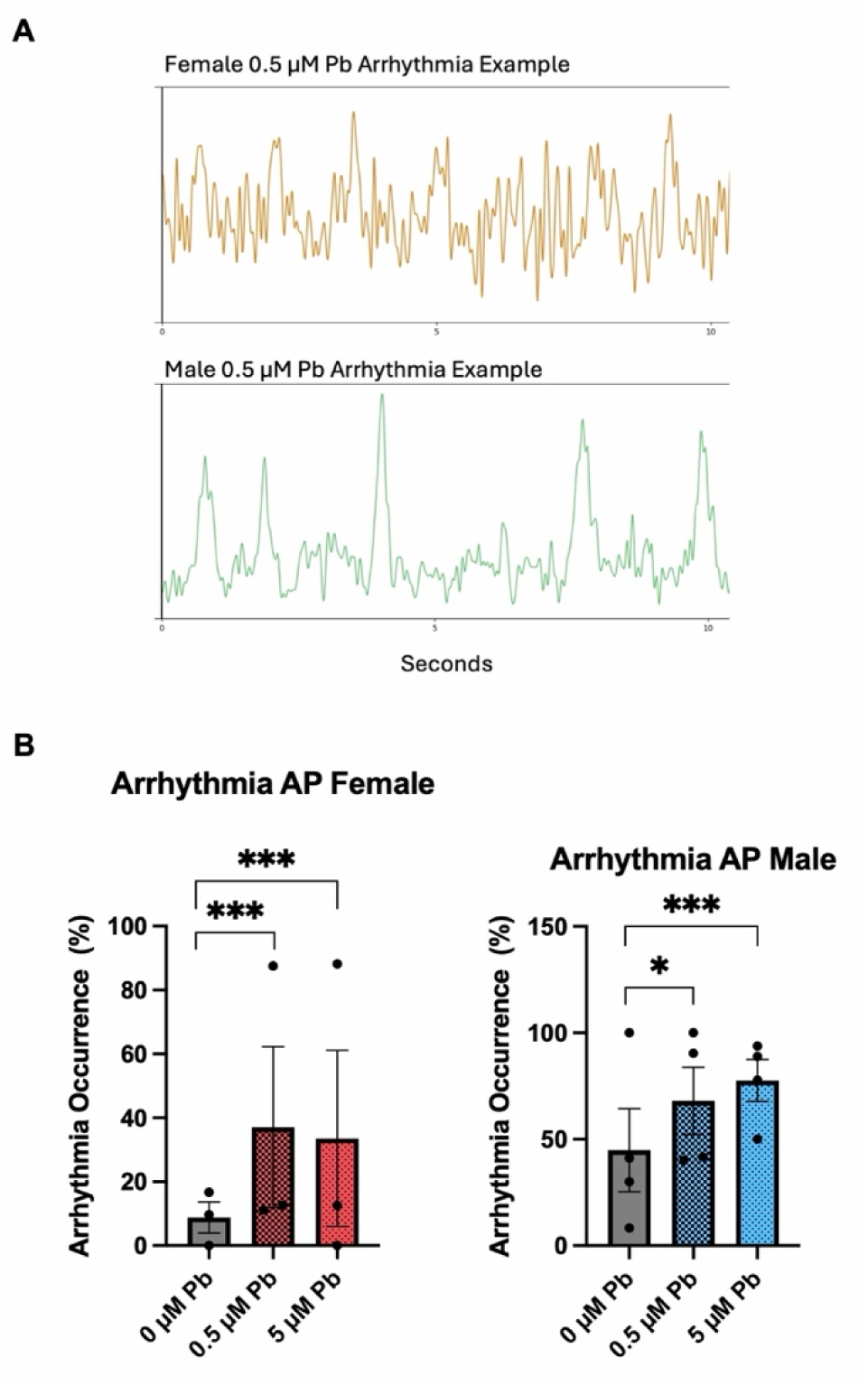
Pb-induced significant increases in action potential percent arrhythmias in female and male iPSC-derived ventricular cardiomyocytes exposed to Pb during differentiation. Percent arrhythmia occurrence was computed by assessing each region’s beat trace for non-consistent patterns (beat-to-beat variation) as demonstrated in beat traces in (A) and arrhythmia occurrence was graphed in (B). Stars indicate significance from a mixed-effects binomial generalized linear modelling accounting for replicate-to-replicate variation; * p-value < 0.05, ** p-value < 0.01, *** p-value < 0.001, **** p-value < 0.0001.

### Exposure to Pb during differentiation affects cardiac Ca transients in a cell line specific manner

We next examined the effects of Pb exposure on Ca dynamics in parallel plates from the same differentiations that underwent mapping of action potential properties. Sample traces for Ca transients are shown in **Figure 4a**. Consistent with the action potential data, Ca transient beat frequency, or the number of contractions per second, was significantly increased at both concentrations of Pb in both cell lines (**Figure 4b**). Ca transient duration triangulation (CaTD Triangulation), which reflects the kinetics of SERCA-mediated reuptake of Ca from the cytosol into the sarcoplasmic reticulum (SR) and the efflux of Ca from the cell via the Na/Ca exchanger. Longer or more triangulated CaTDs suggest slower Ca clearance, whereas shorter or more rectangular transients indicate more efficient Ca cycling. CaTD Triangulation was decreased in the male line at both Pb concentrations (**Figure 4c**). Ca Transient Duration at 80% Recovery (CaTD80), which also reflects the kinetics of Ca clearance, showed a significant increase in the female cell line at both concentrations and decrease in the male cell line at both concentrations (**Figure 4d**). A significant increase was seen in Ca transient amplitude in the female cell line at both Pb concentrations, while a decrease was observed in the male line only at 5 µM. A slight change in baseline fluorescence (increase) was observed only in the female cell line at both concentrations (**Figure 4e and f**).

**Figure 4:**
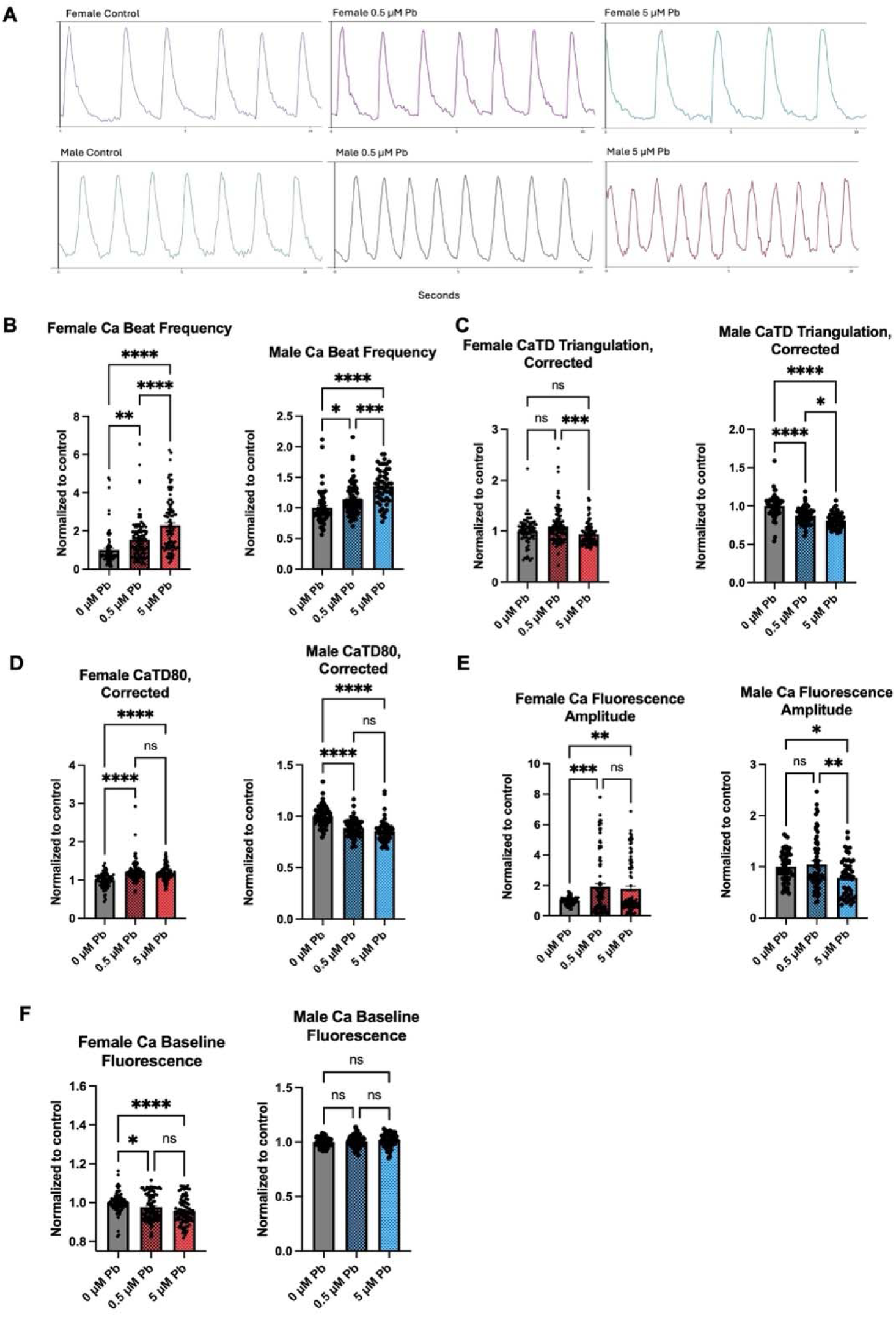
Ca transient changes in female and male iPSC-derived ventricular cardiomyocytes exposed to Pb during differentiation. Sample beat traces for Ca transients shown in (A). Significant changes in beat frequency (B) and CaTD Triangulation (C), CaTD80 (D), Fluorescence Amplitude (E), and Baseline Fluorescence (F) were observed for one or both cell lines. (n= 4 female and n=3 male replicates; one-way ANOVA; * p-value < 0.05, ** p-value < 0.01, *** p-value < 0.001, **** p-value < 0.0001).

As with action potential data, arrhythmias were also assessed for in Ca transient data, but no significant changes were seen for either cell line (**Supplemental Figure 1**).

### No significant Pb-induced ECC gene alterations in iPSC-derived cardiomyocytes

Given the observed Pb-induced changes in action potential and calcium dynamics, which persist long after cessation of exposure, we hypothesized that Pb exposure had reprogrammed expression of genes critical for cardiomyocyte excitability. We therefore examined expression of genes necessary for cardiomyocyte excitation-contraction coupling (ECC) and repolarization using qRT-PCR. Four ECC genes assessed were: *ATP2A2* (SERCA2), which transports Ca from cytosol into sarcoplasmic reticulum, *CACNA1C* (Cav1.2) which transports Ca into the cytosol from the extracellular space, *RYR2* (ryanodine receptor), which transports Ca from sarcoplasmic reticulum into the cytosol, and *SLC8A1* (Na Ca exchanger) which transports Ca out of the cell. *HERG1A* and *B* were also examined, which encode for subunits of a voltage gated potassium (K) channel that mediates repolarizing K currents.(24) However, none of these genes showed any significant expression changes with Pb exposure in purified cardiomyocytes from either cell line (**Figure 5**).

**Figure 5:**
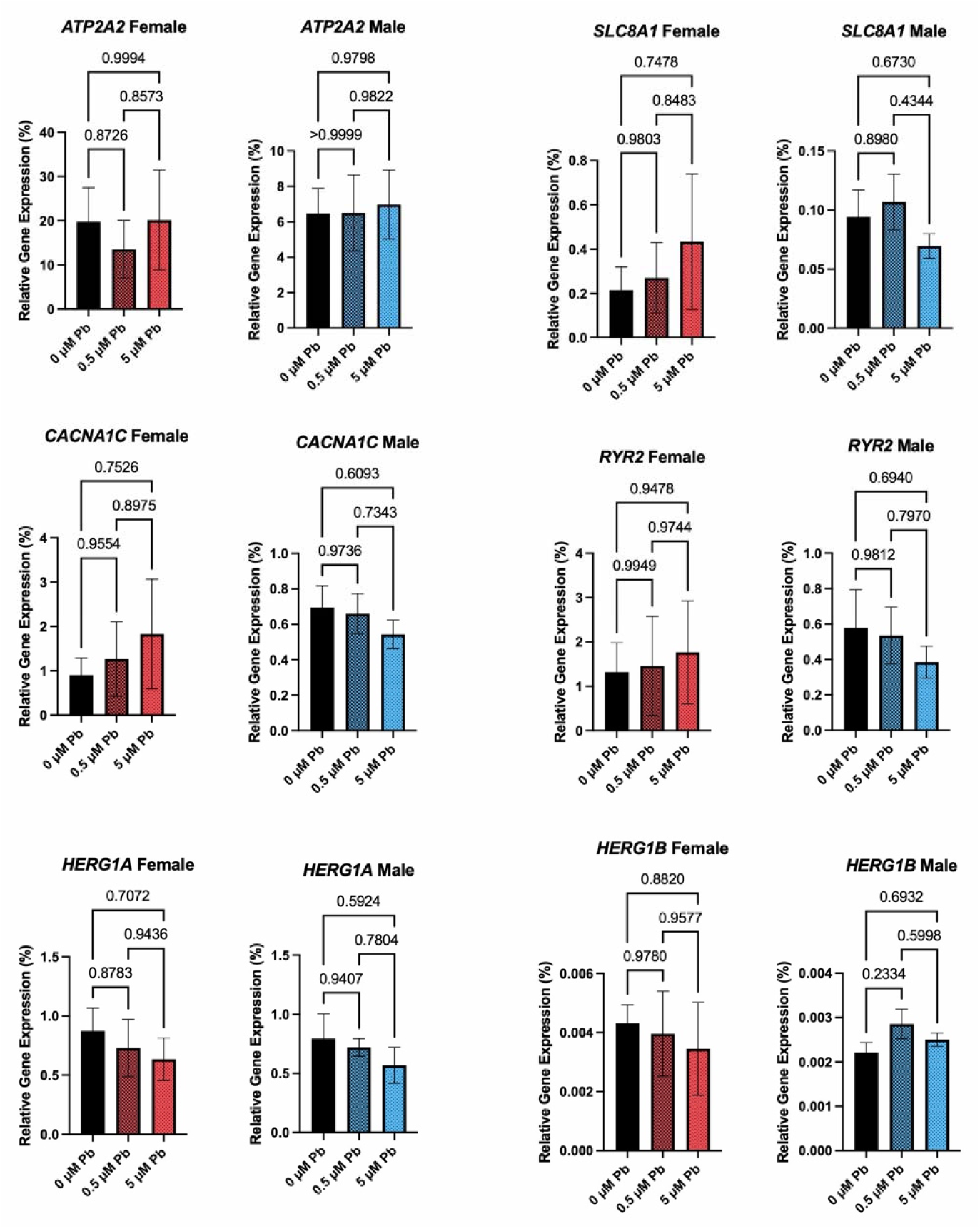
No Pb induced changes in excitation contraction coupling channel/transporter or K channel genes observed for either cell line. For ECC genes *ATP2A2*, *CACNA1C, RYR2*, and *SLC8A1* and K channel subunits *HERG1A* and *B,* no significant changes were observed for purified cardiomyocytes for either female or male cells treated with 0.5 and 5 µM Pb during the first 8 days of differentiation (n = 3 per cell line per treatment; one-way ANOVA p-values for each comparison labeled on the graphs).

## Discussion

Purified cardiomyocytes exposed to Pb (0.5 or 5 µM) during the early stages of differentiation (days 0-8) showed changes in beat characteristics well after exposure had ended, suggesting that Pb exposure can have lasting impacts on human cardiomyocyte function. Interestingly, these effects slightly varied between the two cell lines. To our knowledge, this is the first report of developmental programming by Pb exposure in human cardiomyocytes.

### Pb exposure during differentiation causes cell line-specific changes in action potentials

Effects of Pb on action potential parameters were different between the two cell lines, with an increase in beating rate in the female cell line at 5 µM Pb concentration but a decrease beat rate in male 0.5 µM Pb concentration (**Figure 2b**). At the same time, there was an increase in APD80 in the female cell line and decrease in APD80 in the male cell line with Pb exposure at those same concentrations (**Figure 2c**). Both an increase and decrease in heart rate have the potential to be pro-arrhythmic. Interestingly, in the female line, we observed an increase in heart rate with a simultaneous increase in action potential duration. This prolonged repolarization despite increased firing rate is suggestive of K efflux reduction or delay. In the female cell line, there was also an increase in upstroke slope, or the rate of depolarization, with 0.5 µM Pb, suggesting altered Na channel function. (**Figure 2e**).(25) How Pb impacts K and Na channels in the heart is not fully understood, but studies in hippocampal neurons have shown Pb-induced inhibition of K and Na channels and currents.(26–28) Thus, the observed prolongation of APD80 in the cardiomyocytes may reflect inhibition of K channel function.(26–28) We examined transcript expression of hERG channel, which underlies the repolarizing I_Kr_ current in the heart.(24) Although we observed no changes in expression of *hERG1A* and *B* transcript it is possible that Pb could disrupt either hERG protein expression, relative subunit abundance, posttranslational modifications, and/or function (**Figure 5**). The observed prolongation of action potential duration in the female cell line is notable, given that women tend to have a longer heart rate corrected QT interval and are more prone to drug-induced QT prolongation and arrhythmias, including Torsades de Pointes.(29) Our results thus warrant further investigation using isolated cardiomyocytes from additional female donors.

Notably, in both cell lines, the percent of arrhythmias (defined here as beat irregularities beyond slowed or increased conduction) present increased with both concentrations of Pb, showing that a transient exposure to Pb in a developmental context has arrhythmogenic effects on cardiomyocytes (**Figure 3**). This result is consistent with epidemiological studies which have found that Pb exposure is associated with arrhythmias and conduction abnormalities.(3)

While these results allude to cell line specific disruptions in K and Na currents in cardiomyocytes, little is known about how Pb exposure during differentiation affects early developmental programming of these channels. More studies are needed on the mechanisms by which Pb exposure during differentiation may disrupt the protein expression, regulation, and function of these channels well before cells begin to beat, and how these changes may relate to cardiac arrhythmias and pathology.

### Pb exposure during differentiation causes cell line-specific changes in Ca dynamics

In the male cell line, Pb exposure caused an increase in Ca transient beat frequency with a corresponding decrease in CaTD and CaTD80, indicative of faster Ca handling (**Figure 4b-d**). However, in the female cell line, we found that both beat frequency and CaTD80 increased, indicating potential disruption of Ca efflux via the sodium-calcium exchanger (NCX) or SERCA2a-mediated Ca reuptake into the sarcoplasmic reticulum (**Figure 4b, d).** Pb can interfere with Ca-dependent processes, as they are both divalent cations with similar ionic radii.(30) In both animal models and rodent/human cell models, Pb has been shown to disrupt voltage-gated Ca channels and Ca-dependent neuronal signaling, which may contribute to neurotoxicity.(30–32) In isolated guinea pig hearts, Pb (0-100 µM) causes an increase heart rate, blocks L-type Cav1.2 channels (which allows extracellular Ca into cardiomyocytes), and increases their fast inactivation. Interestingly, L-type Ca channels play a role in embryonic development of the heart and modifications to these channels are associated a variety of arrhythmias such as atrial fibrillation.(33) Pb exposure during differentiation could thus disrupt these channels during development, leading to lasting changes in cardiac electrophysiology. We examined the gene expression of *CACNA1C*, the gene that encodes for the L-type Ca channel and saw no Pb-induced changes in gene expression, suggesting that any disruption likely occurs post-translationally (**Figure 5**).

### Potential for action potential and Ca uncoupling in male ventricular cardiomyocytes exposed to Pb during differentiation

In the female derived cell line, there was an increased beat frequency measured by both action potentials and Ca transients with 5 uM Pb, indicating faster pacing caused by Pb exposure (**Figure 2b** and **4b**). In the male derived cell line, the effects of Pb during differentiation on electrophysiology were more complex. Action potential frequency decreased at the lower concentration, while Ca transient frequency increased at both concentrations (**Figures 2b** and **4b**). This paradoxical shift indicates a slowing of electrical signal propagation with a simultaneous change in Ca kinetics, perhaps originating from an uncoupling of excitation and contraction. Faster Ca transient frequency that is not synchronized with electrical excitation may reflect either compensatory changes in Ca handling machinery within individual cells to maintain contraction despite slowed electrical propagation or maladaptive Ca dysregulation resulting from the slowed electrical stimulus.

Given these possible dysfunctions in ECC, we examined the gene expression of 4 channels/transporters related to ECC in both cell lines and saw no significant gene expression changes (**Figure 5**), Recent work in the heart has found little correlation between transcript and protein changes during murine heart development; thus, future work should include an analysis of the proteome and posttranslational modifications as well.(34) Furthermore, patch clamping of cardiomyocytes exposed to Pb during differentiation should be carried out to examine changes in ion channel function.

### Limitations and future directions

This study is the first to demonstrate how Pb exposure during cardiomyocyte differentiation can disrupt electrophysiology well after exposure has ceased using two independent human cell lines. Importantly, Pb exposure did not affect the timing of when beating first occurred nor the recovery time after purification. It is, however, critical to note that Pb may persist in cells after cessation of exposure, and that could be a source of persistent effects. Small amounts of Pb can accumulate in the heart after exposure, but no work has been done to determine if cardiomyocytes themselves can retain Pb.(35)

There are prominent sex differences in cardiovascular disease, which are likely a function of both hormones and sex chromosomal complement.(36) The cell line-specific differences observed in this study are intriguing, and warrant further investigation using a larger cohort of patient-derived cell lines, or isogenic XX and XY cells to assess the contribution of sex chromosome complement to the observed differential effects of Pb..(37,38)

Additionally, the cardiomyocytes examined here represent a fetal like state, limiting any conclusions related to adult cardiovascular disease development, but by investigating early changes in electrophysiology, we can predict how early life exposures to Pb relate to later cardiac dysfunction or pathology. Future studies currently underway by the lab will examine overall transcriptomic changes in these cells, as well as how ECC and action potential transporters/channels may be post transcriptionally or otherwise functionally regulated by Pb exposure.

## Conclusions

Overall, in this work we characterized electrophysiological changes caused by developmental Pb exposure, revealing persistent and cell line-specific Pb induced effects on beat characteristics in the two cell lines. While no channel or transporter gene expression changes were detected, this study nonetheless demonstrates that early exposure to Pb during cardiomyocyte differentiation can have lasting effects on cardiac electrophysiology.

## Supplementary Information

Supplemental data for this work includes 1 supplemental figure.

## Acknowledgements

I would like to thank the University of Michigan Frankel Cardiovascular Center Cell Core Laboratory, the DoGoodS-Pi Epigenetics Research Group, and the Svoboda lab, including Isabelle Melis, Dr. Tomoko Ishikawa, and Tamara Jones for their laboratory and administrative support. This work was funded by three years of the Environmental Toxicology and Epidemiology (ETEP) training grant (T32 ES0077062) as well as by National Institutes of Health grants: R35 ES031686, P30 ES017885, and K01 ES03204812. Further funding was obtained from the Michigan Biological Research Initiative on Sex Differences in Cardiovascular Disease.

## Author Contributions

Study design was performed by Dr. Kimberley Sala-Hamrick under the guidance of Dr. Laurie Svoboda. Material preparation, data collection and analysis were performed principally by Dr. Kimberley Sala-Hamrick, with aid from Esha Shah. Dr. Andre Monterio provided key expertise in data processing and analysis of optical mapping data. The first draft of the manuscript was written by Dr. Sala-Hamrick. All authors read and approved the final manuscript.

## Supplemental Figures

**Supplemental Figure 1:**
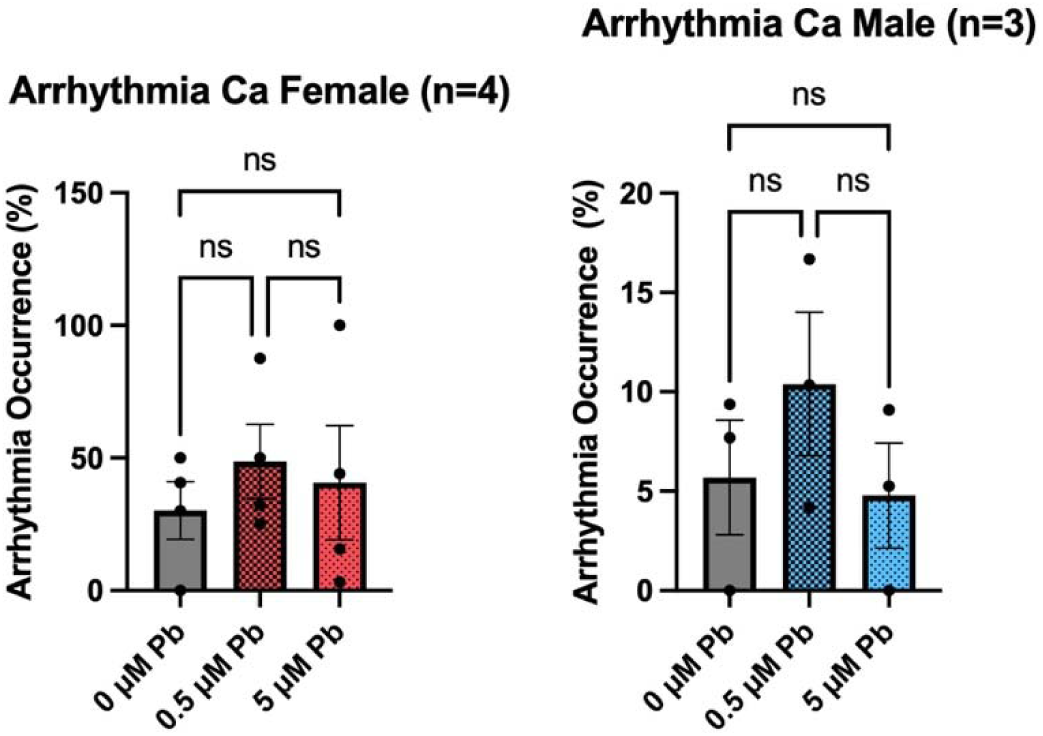
No significant Pb induced changes in Ca transient percent arrhythmias in female and male iPSC-derived ventricular cardiomyocytes treated with Pb during differentiation. Percent arrhythmia occurrence was computed by assessing each region’s beat trace for non-consistent patterns (beat-to-beat variation) and arrhythmia occurrence was graphed. No significant changes were determined using a mixed-effects binomial generalized linear modeling accounting for replicate-to-replicate variation.

